# riboSeed: leveraging prokaryotic genomic architecture to assemble across ribosomal regions

**DOI:** 10.1101/159798

**Authors:** Nicholas R. Waters, Florence Abram, Fiona Brennan, Ashleigh Holmes, Leighton Pritchard

**Affiliations:** Department of Microbiology, School of Natural Sciences, National University of Ireland, Galway, Ireland; Information and Computational Sciences, James Hutton Institute, Invergowrie, Dundee DD2 5DA, Scotland; Soil and Environmental Microbiology, Environmental Research Centre, Teagasc, Johnstown Castle, Wexford, Ireland; Cell and Molecular Sciences, James Hutton Institute, Invergowrie, Dundee DD2 5DA, Scotland

**Keywords:** genome assembly, ribosome, benchmarking, scaffolding, *de fere novo*

## Abstract

The vast majority of bacterial genome sequencing has been performed using Illumina short reads. Because of the inherent difficulty of resolving repeated regions with short reads alone, only ≈10% of sequencing projects have resulted in a closed genome. The most common repeated regions are those coding for ribosomal operons (rDNAs), which occur in a bacterial genome between 1 and 15 times, and are typically used as sequence markers to classify and identify bacteria. Here, we exploit conservation in the genomic context in which rDNAs occur across taxa to improve assembly of these regions relative to *de novo* sequencing by using the conserved nature of rDNAs across taxa and the uniqueness of their flanking regions within a genome. We describe a method to construct targeted pseudocontigs generated by iteratively assembling reads that map to a reference genome’s rDNAs. These pseudocontigs are then used to more accurately assemble the newly-sequenced chromosome. We show that this method, implemented as riboSeed, correctly bridges across adjacent contigs in bacterial genome assembly and, when used in conjunction with other genome polishing tools, can assist in closure of a genome.

## Background

Sequencing bacterial genomes has become much more cost effective and convenient, but the number of complete, closed bacterial genomes remains a small fraction of the total number sequenced (Figure 1). Even with the advent of new technologies for long-read sequencing and improvements to short read platforms, assemblies typically remain in draft status dueto the computational bottleneck of genome closure [5, 33]. Although draft genomes are often of very high quality and suited for many types of analysis, researchers must choose between working with these draft genomes (and the inherent potential loss of data), or spending time and resources polishing the genome with some combination of *in silico* tools, PCR, optical mapping, re-sequencing, or hybrid sequencing [33,46]. Many *in silico* genome finishing tools are available, and we summarise several of these in Table 1. The Illumina entries in NCBI’s Sequence Read Archive (SRA) [22]outnumber all other technologies combined by about an order of magnitude (Table S1). Draft assemblies from these datasets have systematic problems common to short read datasets, including gaps in the scaffolds due to the difficulty of resolving assemblies of repeated regions [44, 53]. By resolving repeated regions during the assembly process, it may be possible to improve existing assemblies, and therefore obtain additional sequence information from existing short read datasets in the SRA or the European Nucleotide Archive.

**Figure 1:**
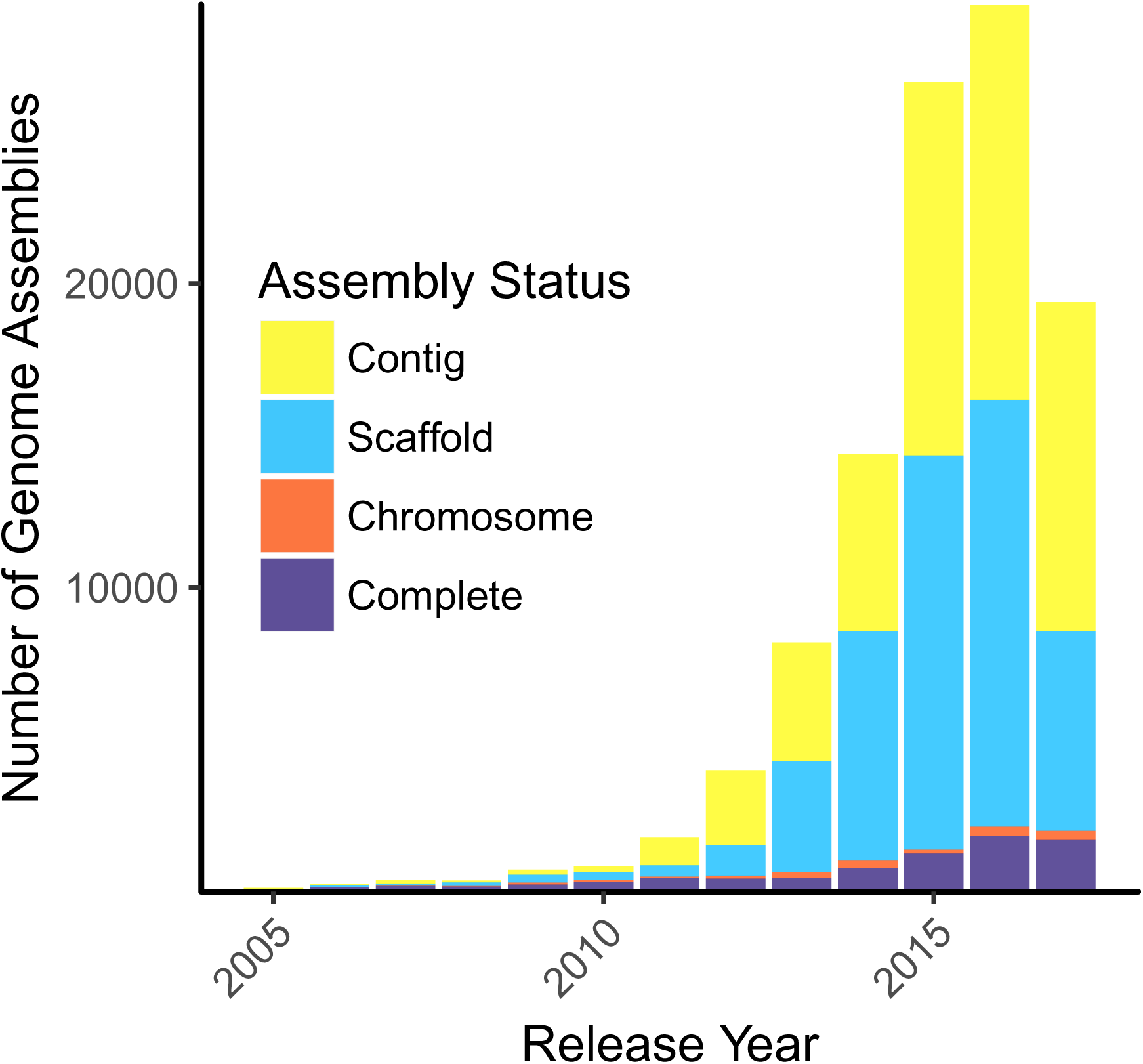
Counts of bacterial assemblies in NCBI Genome database according to completion level by release year; the four levels (Complete, Chromosome, Scaffold, and Contig) are ordered from complete to most fragmented [21]. Note that Illumina HiSeq was released in 2010. Accessed September 14^th^, 2017.

**Table 1:**
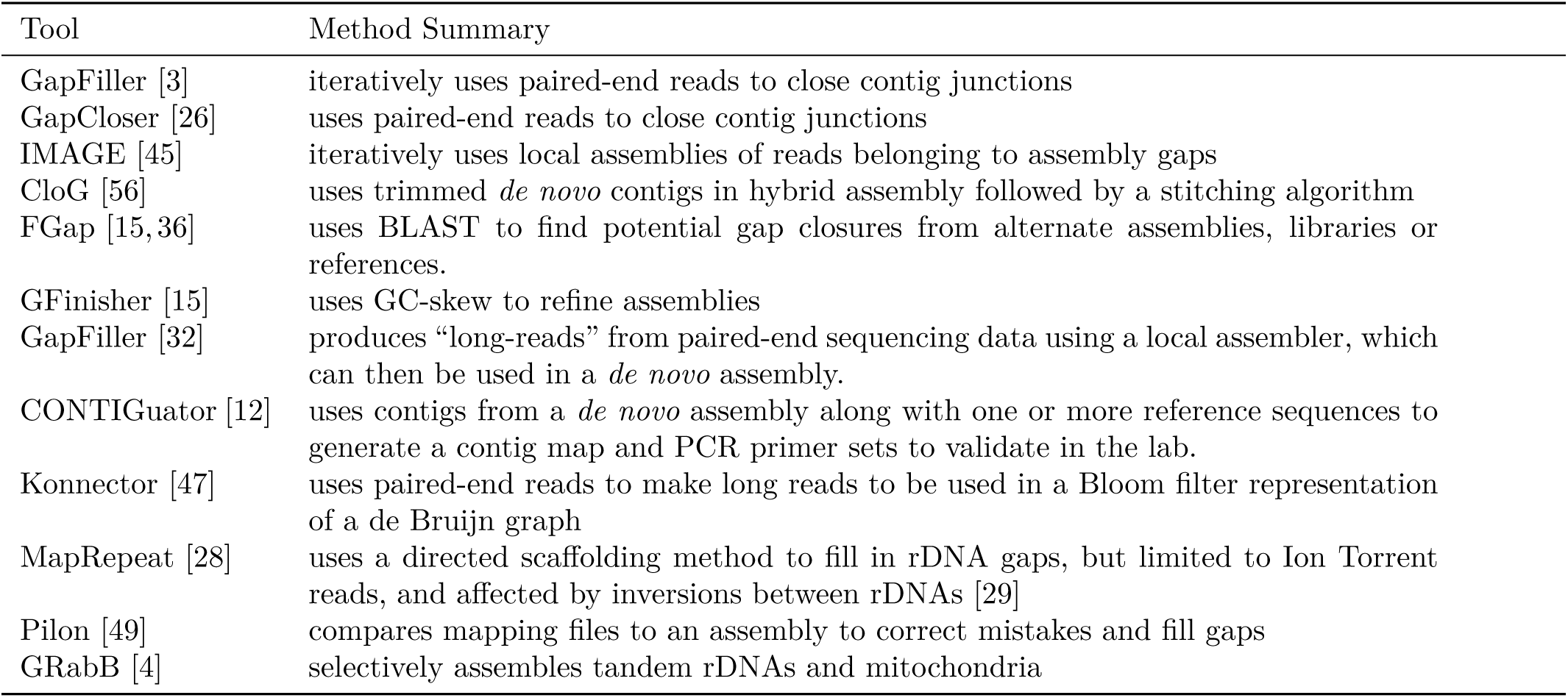
Some of the available in silico genome polishing tools for gap closure

The most common repeated regions are those coding for ribosomal RNA operons (rDNAs). As ribosomes are essential for cell function, sequencing of the 16S ribosomal region is widely used to identify prokaryotes and explore microbial community dynamics [6, 7, 52, 54]. This region is conserved within taxa, yet retains enough variability to act as a bacterial “fingerprint” to separate clades informatively. However, the 16S, 23S, and 5S ribosomal subunit coding regions are often present in multiple copies within a single prokaryotic genome, and commonly exhibit polymorphism [8,25,31,48]. These long, inexactly repeated regions [1] are problematic for short-read genome assembly. As rDNAs are frequently used as a sequence marker for taxonomic classification, resolving their copy number and sequence diversity from short read collections where the assembled genome has collapsed several repeats into a single region could help improve reference databases, increasing the accuracy of community analysis. We present here an *in silico* method, riboSeed, that capitalizes on the genomic conservation of rDNA and flanking sequence within a taxon to improve resolution of these difficult regions and provide a means to benefit from unexploited information in the SRA/ENA short read archives.

riboSeed is most similar in concept to GRabB, the method of Brankovics et al. [4] for assembling mitochondrial and rDNA regions in eukaryotes. Both use targeted assembly, but GRabB does not make inferences about the number of rDNA clusters present in the genome, or take advantage of their genomic context. In riboSeed, genomic context is resolved by exploiting both the rDNA sequences and their flanking regions, harnessing unique characteristics of the broader rDNA region within a single genome to improve assembly.

The riboSeed algorithm proceeds from two observations: first, that although repeated rRNA coding sequences within a single genome are nearly identical, their flanking regions (that is, the neighboring locations within the genome) are distinct in that genome, and second, that the genomic contexts of equivalent rDNA sequences are also conserved within a taxonomic grouping (Figure S4). riboSeed uses only reads that map to rDNA regions from a reference genome, and is not affected by chromosomal rearrangements that occur outside the flanking regions immediately adjacent to each rRNA.

Briefly, riboSeed uses rDNA regions from a closely-related organism’s genome to help generate rDNA cluster-specific “pseudocontigs” derived only from the input short reads, that are seeded into the raw short reads to generate a final assembly. We refer to this process in this work as *de fere novo* (meaning “starting from almost nothing”) assembly.

### Implementation

We present riboSeed: a software suite that allows users to perform *de fere novo* assembly, given a reference genome sequence from a closely-related organism and single or paired-end short reads to be assembled (Figure 2). The code is primarily written in Python3, with accessory shell and R scripts. riboSeed relies on a closed reference genome assembly that is sufficiently closely-related to the isolate being assembled (distance can be estimated using an alignment-free approach such as the KGCAK database [50], or a kmer based method such as Kraken [55]), in which rDNA regions are assembled and assumed to be in the correct genomic context.

**Figure 2:**
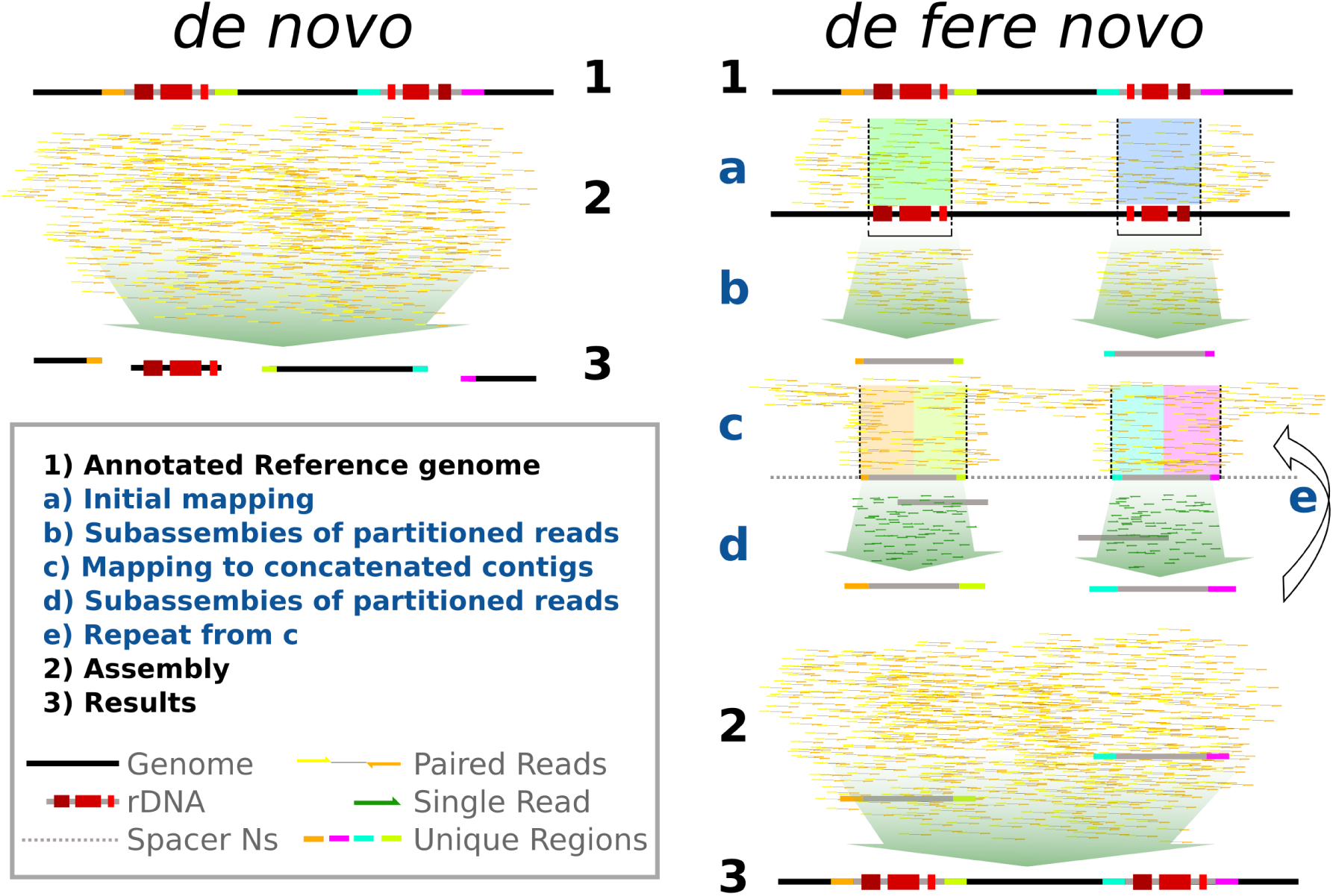
A comparison of *de novo* assembly to *de fere novo* assembly, as implemented in riboSeed. In riboSeed, reads are mapped to a reference genome, and those reads that align to rDNA and flanking regions are extracted. A subassembly for each group of reads that maps to an rDNA region is constructed to produce a “pseudocontig” for each region. These pseudocontigs are concatenated together separated by 1kb of Ns as a spacer. Reads are then iteratively mapped to the concatenated pseudocontigs, extracted, and again subassembled to each region. After the final iteration, the pseudocontigs are included with raw reads in a standard *de novo* assembly. The subassemblies attempt to bridge proper rDNA regions by ensuring that flanking regions (represented here by colors) remain correctly paired. The *de novo* assembly collapses the rDNAs, but *de fere novo* places the rDNAs in the proper genomic context.

In an ideal scenario, reference selection would consist of two steps: isolate identification (using Kraken), and then average nucleotide identity analysis to find the closest complete reference. We outline protocols for reference selection in Supplementary Information, and in the riboSeed documentation at http://riboseed.readthedocs.io/en/latest/REFERENCE.html.

### Usage

Installation (via either conda, pip, or GitHub), installs the `ribo` program. Installation using conda also installs third-party tool dependencies, such as SPAdes, and is recommended. The riboSeed pipeline can be executed with a single command, `ribo run`, or under stepwise control by the user by means of distinct commands. `ribo run` performs reannotation of rDNAs in the reference genome (`scan` command), operon inference (`select` command), and *de fere novo* assembly (`seed` command). The most commonly used parameters are accessible via the `run` command. Alternatively, the full set of parameters for riboSeed can be defined within a configuration file.

All steps in the assembly are controlled by riboSeed commands described below, as `ribo <command>:`

#### 1. Preprocessing

##### scan

`scan` uses Barrnap (https://github.com/tseemann/barrnap) to annotate rRNAs in the reference genome, and EM-BOSS’s seqret [38] to create GenBank, FASTA, and GFF formatted versions of the reference genome. This preprocessing step unifies the annotation vocabulary for downstream processes.

##### select

`select` infers ribosomal operon structure from the genomic location of constituent 16S, 23S and 5S sequences. Jenks Natural Breaks algorithm is used to group rRNA annotations into likely operons on the basis of genomic coordinates, using the number of 16S annotations to set the number of breaks. Output defines individual rDNA clusters and describes component elements in a plain text file. This output can be manually adjusted before assembly if clustering does not reflect the known arrangement of operons, for example based on visualization of the annotations in a genome browser.

#### 2. De Fere Novo Assembly

##### seed

`seed` implements the algorithm described in Figure S1. Short reads for the sequenced isolate are mapped to the reference genome using BWA [23]. Reads that map to each annotated rDNA and its flanking regions (where the flanking regions consist of 1kb upstream and 1kb downstream of the rDNA, by default) are extracted into subsets (one subset per cluster). Each subset is independently assembled into a representative pseudocontig with SPAdes [2], using the reference rDNA regions as a trusted contig (or untrusted, if mapping quality is poor). Resulting pseudocontigs are evaluated for inclusion in future mapping/subassembly iterations based on length, and concatenated into a pseudogenome in which pseudocontigs are separated by 1kb of Ns as a spacer. As we are only concerned with flanking regions, the order in which the pseudocontigs are concatenated is arbitrary. A 1kb spacer length was chosen for this study to ensure that reads did not span the spacer. Pseudocontig generation is repeated in each iteration of the algorithm, using the previous round’s pseudogenome as the reference.

After a specified number of iterations (3 by default), SPAdes is used to assemble all short reads in a hybrid assembly using pseudocontigs from the final iteration as “trusted contigs” (or “untrusted contigs” if the mapping quality of reads to that pseudocontig falls below a threshold). As a control, the short reads are also *de novo* assembled without the pseudocontigs.

This implementation of riboSeed uses SPAdes to perform both subassembly and the final *de fere novo* assembly, but the pseudocontigs could be submitted to any hybrid assembler that accepts short read libraries and contigs. After assembly, *de fere novo* and *de novo* assemblies are assessed with QUAST [16].

#### 3. Assessment and Visualization

##### score

`score` extracts the regions flanking rDNAs in the reference and in assemblies generated by riboSeed. Flanking regions from an assembly are matched with reference flanking regions using BLASTn. Depending on the ordering of the matches, assembled junctions are called as correct, incorrect, or ambiguous based on the criteria outlined below.

##### snag

`snag` is a helper tool to produce diagnostics and visualisation of rDNA sequences in the reference genome. Using the clustering generated by `select`, sequences for the clusters are extracted from the genome, aligned, and Shannon entropy [41] plotted with consensus depth for each position in the alignment.

##### swap

We recommend assessing the performance of the riboSeed pipeline visually using Mauve [9, 10], Gingr [43], or similar genome assembly visualizer to compare reference, *de novo,* and *de fere novo* assemblies. If contigs appear to be incorrectly joined, the offending *de fere novo* contig can be replaced with syntenic contigs from the *de novo* assembly using the `swap` script.

##### stack

`stack` uses bedtools [37] and samtools [23] to compare depth of coverage of reads aligning to the reference genome in the rDNA regions to randomly sampled regions elsewhere in the reference genome. `stack` takes output from `scan`, and a BAM file of reads that map to the reference. If the number of `scan`-annotated rDNAs matches the number of rDNAs in the sequenced isolate, the coverage depths within the rDNAs will be similar to the coverage in other locations in the genome. If the coverage of rDNA regions sufficiently exceeds the average coverage elsewhere in the genome, this may indicate that the reference strain has fewer rDNAs than the sequenced isolate. In this case, using an alternative reference genome may produce improved results.

### Availability of data and materials

The riboSeed pipeline and datasets generated in this study are available on the riboSeed website, https://nickp60.github.io/riboSeed/. The software is released under the MIT licence. The modified BugBuilder pipeline used here is provided at https://github.com/nickp60/BugBuilder. Reference genomes used for this study can be found in Table S3, and the versions of other software used in this study are found in Table S4.

#### Choice of parameters

Settings used for analyses in this manuscript are default (except where otherwise noted) as of riboSeed version 0.4.35 [51].

#### Validating assembly across rDNA regions

To evaluate performance of *de fere novo* assembly compared to *de novo* assembly methods, we used Mauve to visualize syntenic regions and contig breaks of each riboSeed assembly in relation to the reference genome used to generate pseudocontigs. We categorized each rDNA in an assembly as either correct, unassembled, incorrect, or ambiguous, as follows.

An rDNA assembly is classed as “correct” if two criteria are met: (i) the assembly joins two contigs across an rDNA region such that, based on the reference, the flanking regions of the *de fere novo* assembly are syntenous with those of the reference; and (ii) the assembled contig extends at least 90% of the flanking region length. A cluster is defined as “unassembled” if the ends of one or more contigs align within the rDNA or flanking regions (extension across the rDNA region is not achieved). Finally, if two contigs assemble across a rDNA region in a manner that conflicts with the orientation indicated in the reference genome, suggesting missassembly, the rDNA region is classified as “incorrect”.

For analyses where manual inspection was intractable (such as repeated simulations), `ribo score` was used to categorize the rDNA assemblies. In cases where the program could not distinguish between a correct assembly or an incorrect assembly, the rDNA was classed as “ambiguous”.

In all cases, SPAdes was used with the same parameters for both *de fere novo* assembly and *de novo* assembly, apart from addition of pseudocontigs in the *de fere novo* assembly.

## Results

### Characteristics of rDNA flanking regions

The use of rDNA flanking sequences to uniquely identify and place rDNAs in their genomic context requires their flanking sequences to be distinct within the genome for each region. This is expected to be the case for nearly all prokaryotic genomes where rRNA coding sequences are structured as operons. We determined that a 1kb flanking region was sufficient to include differentiating sequence (Figure S2). To demonstrate this, rDNA and 1kb flanking regions were extracted from *E. coli* Sakai [17] (BA000007.2), a strain in which rDNAs have been well characterized [34]. These regions were aligned with MAFFT [20], and consensus depth and Shannon entropy calculated for each position in the alignment (Figure 3a).

**Figure 3:**
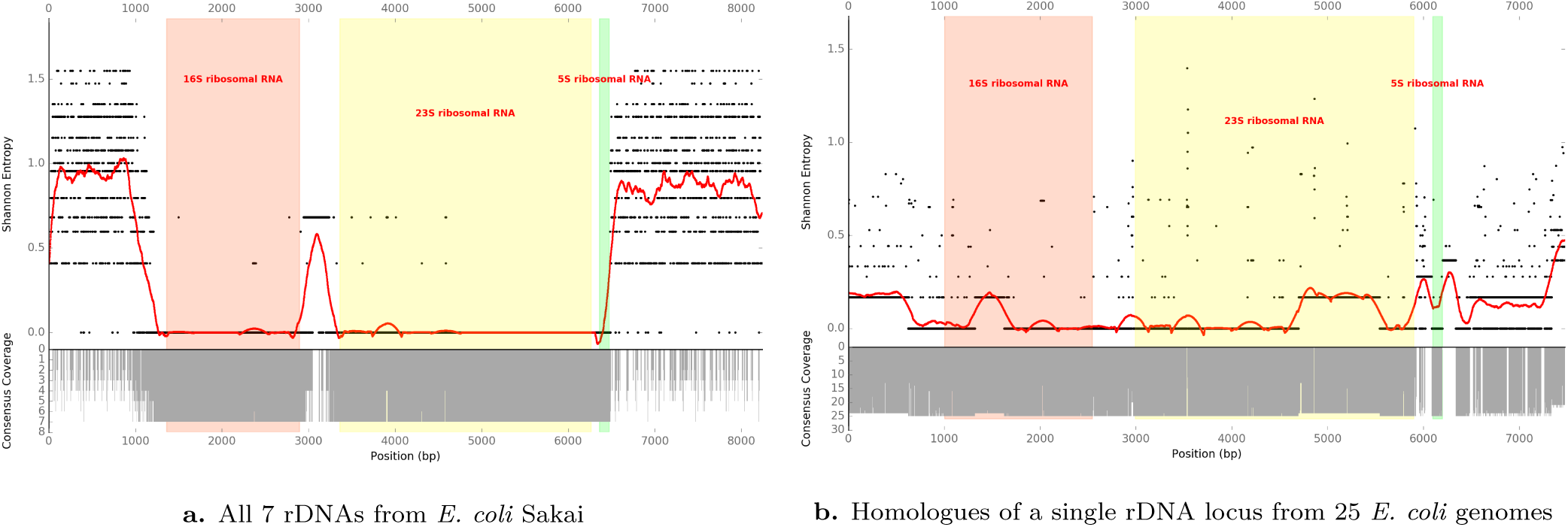
Consensus coverage depth (gray bars) and Shannon entropy (black points, smoothed with a window size of 351bp as red line) for aligned rDNA regions. For the seven *E. coli* Sakai rDNA regions (a), entropy sharply increases moving away from the 16S and 5S ends of the operon. In this case flanking regions would be expected to assemble uniquely within a genome. By contrast, the rDNA regions occurring closest to homologous *gmhB* genes from 25 *E. coli* genomes (b) show greater conservation in their flanking regions. This indicates that flanking regions are more conserved for homologous rDNA than for paralogous rDNA operons, and implies that related genomes can be useful reference templates for assembling across these regions. Similar plots for each of the GAGE-B genomes used later for benchmarking can be found in Figure S4.

Figures 3a and S4 show that within a single genome the regions flanking rDNAs are variable between operons. This enables unique placement of reads at the edges of rDNA coding sequences in their genomic context (i.e. there is not likely to be confusion between the placements of rDNA edges within a single genome). In *E. coli* MG1655 (NC_000913.3), the first rDNA is located 363 bases downstream of *gmhB* (locus tag b0200). Homologous rDNA regions were extracted from 25 randomly selected complete *E. coli* chromosomes (Table S2). We identified the 20kb region surrounding *gmhB* in each of these genomes, then annotated and extracted the corresponding rDNA and flanking sequences. These sequences were aligned with MAFFT, and the Shannon entropies and consensus depth plotted (Figure 3b).

Figure 3b shows that equivalent *E. coli* rDNAs, plus their flanking regions, are well-conserved across several related genomes. Assuming that individual rDNAs are monophyletic within a taxonomic group, short reads that can be uniquely placed on a related genome’s rDNA as a reference template are also likely able to be uniquely-placed in the appropriate homologous rDNA of the genome to be assembled.

Taken together, when these two properties hold, this allows for unique placement of reads from homologous rDNA regions in the appropriate genomic context. These “anchor points” effectively reduce the number of branching possibilities in de Bruijn graph assembly for each individual rDNA, and thereby permit reconstruction of a complete balanced path through the full rDNA region.

### Simulated reads with artificial chromosome

To create a small dataset for testing, we extracted all 7 distinct rDNA regions from the *E. coli* Sakai genome (BA000007.2), including 5kb upstream and downstream flanking sequence, using the tools `scan`, `select` and `snag`. Those regions were concatenated to produce a ~100kb artificial test chromosome (see supplementary methods). pIRS [19] was used to generate simulated reads (100bp, 300bp inserts, stdev 10, 30-fold coverage, built-in error profile) from this test chromosome. These reads were assembled using riboSeed, using the *E. coli* MG1655 genome (NC_000913.3) as a reference. Simulation was repeated 8 times to assess variability of method performance on alternative read sets generated from the same source sequence; Figure 4 shows a Mauve visualization of a representative run. *de fere novo* assembly bridged 4 of the 7 rDNA regions in the artificial chromosome, while *de novo* assembly failed to bridge any (Figure S3). To illustrate how choice of reference sequence determines correct assembly through rDNA, we ran riboSeed with the same *E. coli* reads using pseudocontigs derived from the *Klebsiella pneumoniae* HS11286 (CP003200.1) reference genome [24]. *de fere novo* assembly with pseudocontigs from *K. pneumoniae* failed, as the reference is too divergent from the reads.

**Figure 4:**
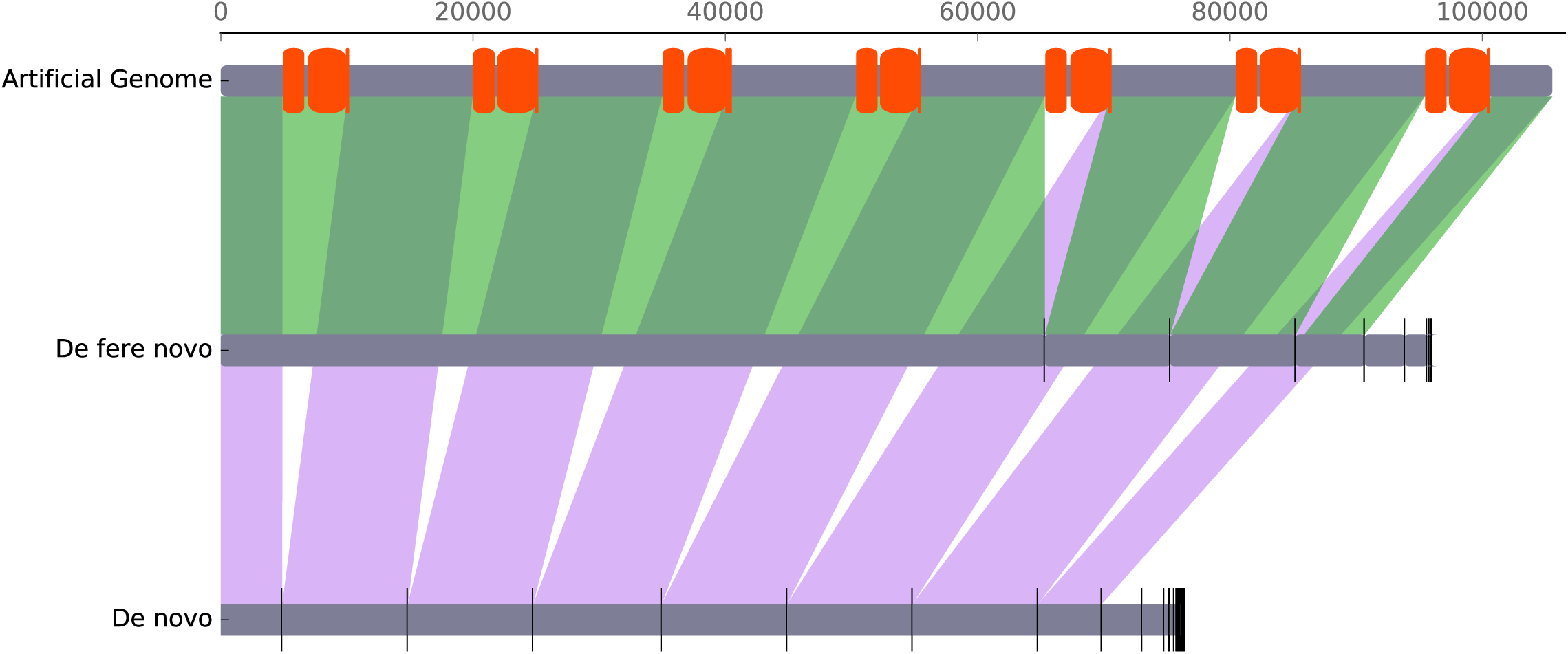
Representative Mauve output describing the results of riboSeed assemblies of simulated reads generated by pIRS from the concatenated *E. coli* Sakai artificial chromosome. Red regions represent rRNA coding sequences, vertical black lines indicate boundaries between assembled contigs, and shading represents synteny. From top to bottom: artificial reference chromosome; rDNA clusters (red bars); *de fere novo* assembly and *de novo* assembly (both using *E. coli* MG1655 as the reference). riboSeed’s *de fere novo* method assembles 4 of 7 rDNA regions, but the *de novo* assembly recovers no rDNA regions correctly.

### Effect of reference sequence identity on riboSeed performance

To investigate how riboSeed assembly is affected by choice of reference strain, we implemented a simple mutation model to generate reference sequence variants of the artificial chromosome described above, with a specified rate of mutation. A simple model of geometrically-distributed mutations at a desired mutation frequency applied across all bases uniformly does not address the disparity of conservation between rDNAs and their flanking region observed in nature, so a second model was applied wherein substitutions are restricted to the rDNA flanking regions. We assembled the artificial chromosome’s reads using the mutated artificial chromosome as a reference, using both models (Figure 5). The maximum substitution rate exceeded our recommended threshold sequence identity, and a corresponding dropoff of performance is observed at a value of 0.2 (corresponding to the 80% mapping percentage identity threshold). To obtain an estimate of substitution rate for the *E. coli* strains used above, Parsnp [43] and Gingr [43] were used to identify SNPs in the 25 genomes used in Figure 3, with respect to the same region in *E. coli* Sakai. An average substitution rate of ≈ 3.5 substitutions per kb was observed. Compared to the results from simulated genomes, we expect successful riboSeed performance under the model of mutated flanking regions, and partial success under the model of substitutions throughout the region.

**Figure 5:**
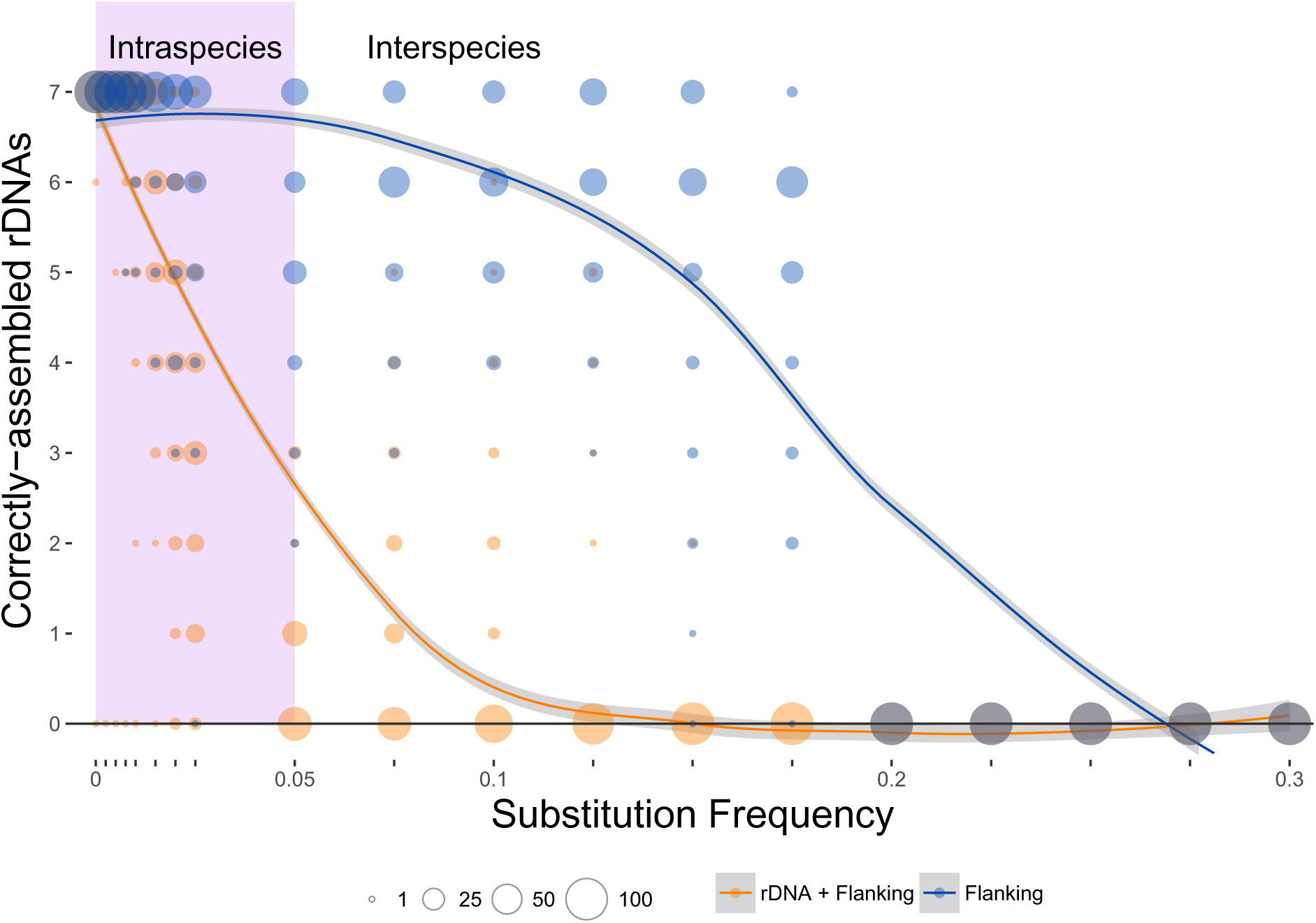
Variants of the artificial chromosome with substitution frequencies between 0 and 0.3 (i.e. up to 300 substitutions per kb). Correctly-assembled rDNAs were counted, and the distribution of results shown against the appropriate substitution frequency. Results are shown for models where substitutions are permitted throughout the chromosome (orange), and only in the flanking regions (blue), the latter approximating the relative rate of substitution in rDNA and flanking regions. The lilac area corresponds to substitution frequencies resulting in average sequence identity over 95%, denoting an estimated species boundary. Loess smoothing was used to generate the blue and yellow trendlines. Circle size indicates number of simulations per value. n=100.

Figure 5 indicates that the greater the similarity of the reference sequence to the genome being assembled, the greater the likelihood of correctly assembling all rDNA regions. When mutating only flanking regions (Figure 5), which more closely resembles the relative sustitution frequencies of the rDNA regions, the procedure correctly assembles rDNAs with tolerance to substitution frequencies up to approximately 30 substitutions per kb. With the widely-adopted average nucleotide identity species boundary of 95% [13], we anticipate that riboSeed should correctly place and assemble most rDNA regions when using a complete reference genome of the same species, and that reasonable success will be achieved even when using a more distantly-related reference.

### Simulated reads with *E. coli* and *K. pneumoniae* genomes

To investigate the effect of short read length on riboSeed assembly, pIRS [19] was used to generate paired-end reads from the complete *E. coli* MG1655 and *K. pneumoniae* NTUH-K2044 genomes, simulating datasets at a range of read lengths most appropriate to the sequencing technology. In all cases, 300bp inserts with 10bp standard deviation and the built-in error profile were used. Coverage was simulated at 20x to emulate low coverage runs and at 50x to emulate coverage close to the optimized values determined by Miyamoto [30] and Desai [11]. *De fere novo* assembly was performed with riboSeed using *E. coli* Sakai and *K. pneumoniae* HS11286 as references, respectively, and the results were scored with `Score` (Figure 6).

**Figure 6:**
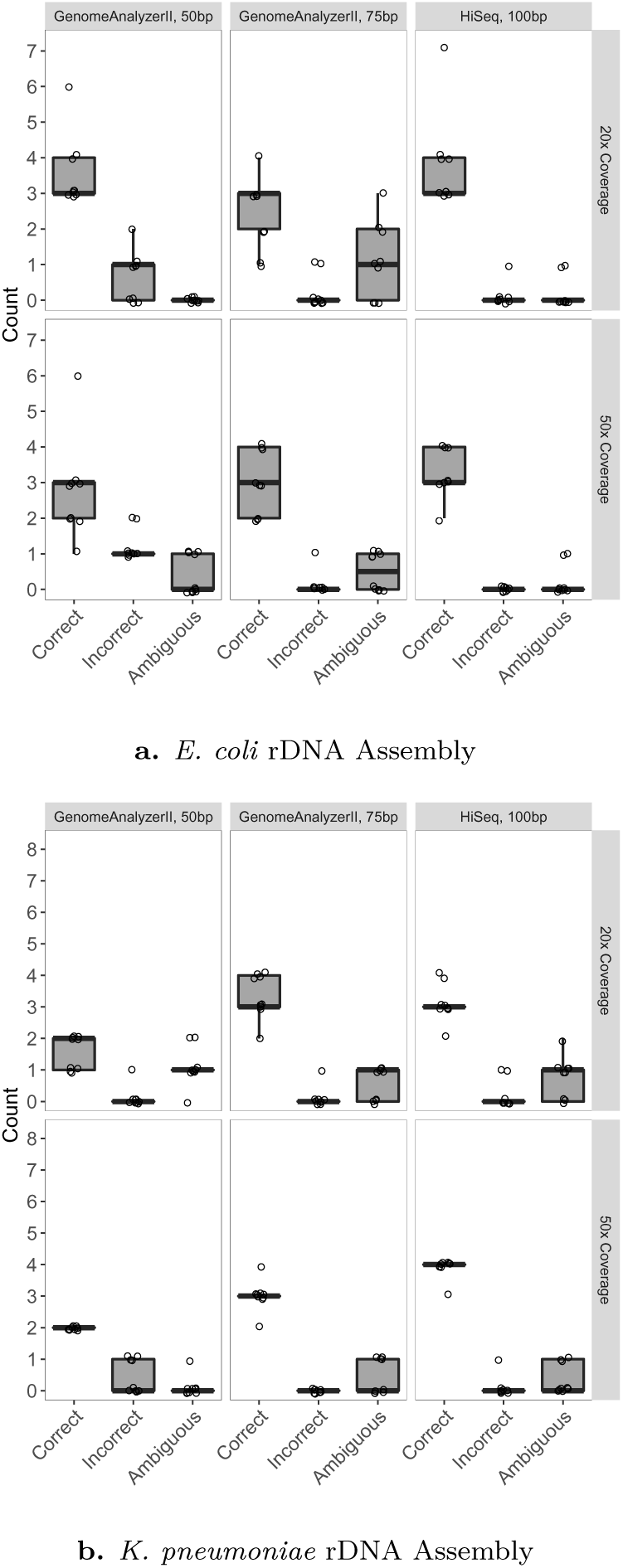
Comparison of *de fere novo* assemblies of simulated reads generated by pIRS. In most cases, increasing coverage depth and read length resulted in fewer misassemblies. Assemblies were scored using `score`; the y axis reflects the total number of rDNAs in the genome (7 and 8 rDNAs, for *E. coli* and *K. pneumoniae* respectively). The (n=9) assemblies shown for each genome are the result of differently seeded read simulations.

At either 20x or 50x coverage, *de novo* assembly was unable to resolve any rDNAs with any of the simulated read sets. *de fere novo* assembly with riboSeed showed improvement to both the *E. coli* and *K. pneumoniae* assemblies. Increasing depth of coverage and read length improves rDNA assemblies.

### Benchmarking against hybrid sequencing and assembly

To establish whether riboSeed performs as well with short reads obtained by sequencing a complete prokaryotic chromosome as with simulated reads, we attempted to assemble short reads from a published hybrid Illumina/PacBio sequencing project. The hybrid assembly using long reads was able to resolve rDNAs directly, and provides a benchmark against which to assess riboSeed performance in terms of: (i) bridging sequence correctly across rDNAs, and (ii) assembling rDNA sequence accurately within each cluster.

Sanjar, *et al.* published the genome sequence of *Pseudomonas aeruginosa* BAMCPA07-48 (CP015377.1) [39], assembled from two libraries: ca. 270bp fragmented genomic DNA with 100bp paired-end reads sequenced on an Illumina HiSeq 4000 (SRR3500543), and long reads from PacBio RS II. The authors obtained a closed genome sequence by hybrid assembly. We ran the riboSeed pipeline on only the HiSeq dataset in order to compare *de fere novo* assembly to the hybrid assembly and *de novo* assembly of the same reads, using the related genome *P. aeruginosa* ATCC 15692 (NZ_CP017149.1) as a reference.

*de fere novo* assembly correctly assembled across all 4 rDNA regions, whereas *de novo* assembly failed to assemble any rDNA regions (Table 3A).

**Table 3:**
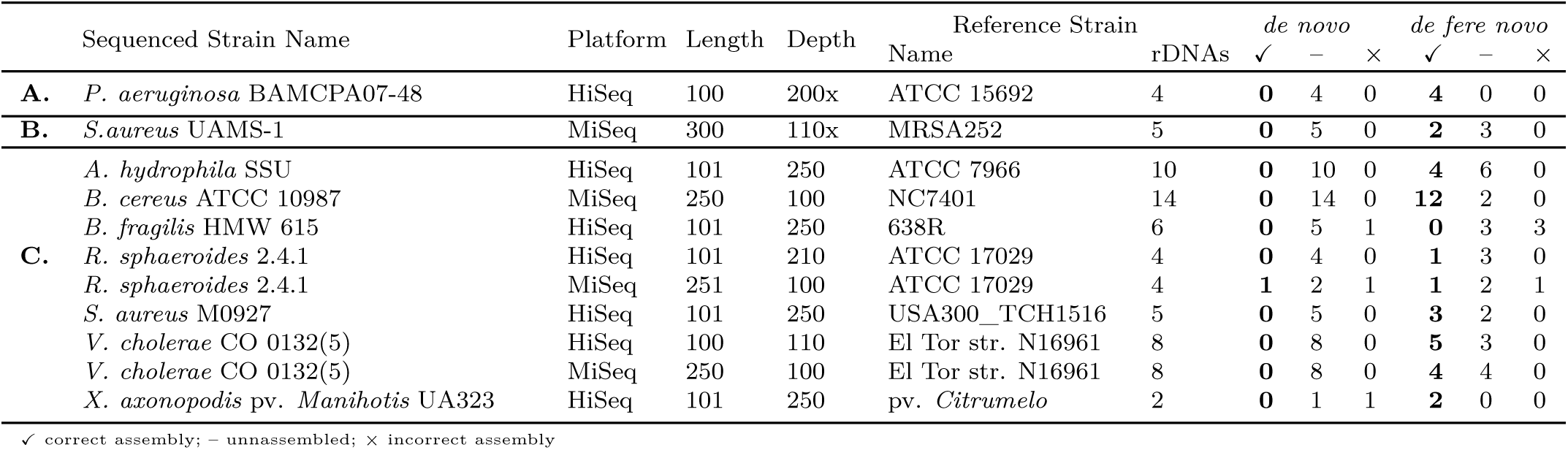
Comparison of *de novo* and riboSeed’s *de fere novo* assemblies

|  | Sequenced Strain Name | Platform | Length | Depth | Reference Strain Name | rDNAs | *de novo* |  |  | *de fere novo* |  |  |
| --- | --- | --- | --- | --- | --- | --- | --- | --- | --- | --- | --- | --- |
|  |  |  |  |  |  |  | ✓ | – | × | ✓ | – | × |
| **A.** | *P. aeruginosa* BAMCPA07-48 | HiSeq | 100 | 200x | ATCC 15692 | 4 | **0** | 4 | 0 | **4** | 0 | 0 |
| **B.** | *S.aureus* UAMS-1 | MiSeq | 300 | 110x | MRSA252 | 5 | **0** | 5 | 0 | **2** | 3 | 0 |
| **C.** | *A. hydrophila* SSU | HiSeq | 101 | 250 | ATCC 7966 | 10 | **0** | 10 | 0 | **4** | 6 | 0 |
|  | *B. cereus* ATCC 10987 | MiSeq | 250 | 100 | NC7401 | 14 | **0** | 14 | 0 | **12** | 2 | 0 |
|  | *B. fragilis* HMW 615 | HiSeq | 101 | 250 | 638R | 6 | **0** | 5 | 1 | **0** | 3 | 3 |
|  | *R. sphaeroides* 2.4.1 | HiSeq | 101 | 210 | ATCC 17029 | 4 | **0** | 4 | 0 | **1** | 3 | 0 |
|  | *R. sphaeroides* 2.4.1 | MiSeq | 251 | 100 | ATCC 17029 | 4 | **1** | 2 | 1 | **1** | 2 | 1 |
|  | *S. aureus* M0927 | HiSeq | 101 | 250 | USA300_TCH1516 | 5 | **0** | 5 | 0 | **3** | 2 | 0 |
|  | *V. cholerae* CO 0132(5) | HiSeq | 100 | 110 | El Tor str. N16961 | 8 | **0** | 8 | 0 | **5** | 3 | 0 |
|  | *V. cholerae* CO 0132(5) | MiSeq | 250 | 100 | El Tor str. N16961 | 8 | **0** | 8 | 0 | **4** | 4 | 0 |
|  | *X. axonopodis* pv. *Manihotis* UA323 | HiSeq | 101 | 250 | pv. *Citrumelo* | 2 | **0** | 1 | 1 | **2** | 0 | 0 |

✓ correct assembly; – unassembled; × incorrect assembly

Comparing the BAMCPA07-48 reference to the *de fere novo* assembly, we found a total of 9 SNPs in the rDNA flanking regions (Table 2). The same regions from the ATCC 15692 reference used in the *de fere novo* assembly showed 108 SNPs compared to the BAMCPA07-48 isolate. This demonstrates that this subassembly scheme successfuly recovers the correct sequence with remarkably few SNPs, despite a large number of differences between the reference and the sequenced isolate, and that the riboSeed method does not simply transpose the reference genome rDNA sequence into the new assembly.

**Table 2:**
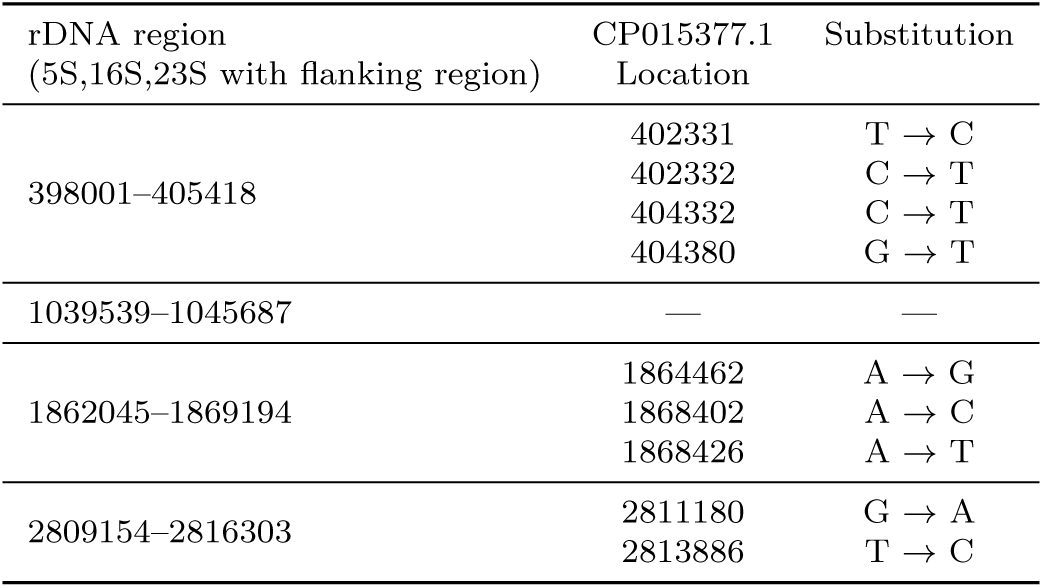
rDNA region SNPs between hybrid assembly of *P. aeruginosa* BAMCPA07-48 and *de fere novo* assembly in rDNA regions, including 1kb upstream and downstream of the rDNA

Further, to assess how riboSeed’s assembly would compare to supplying the whole reference as a trusted contig in SPAdes (a strategy not recommended by the SPAdes authors), we assembled the same reads with the *P. aeruginosa* ATCC 15692 as a trusted contig, and compared results to *de fere novo* and *de novo* assemblies. *de novo* assembly yielded the lowest error rates, and *reference-based* assembly yielded the longest contigs, but *de fere novo* assembly exhibited very low error rates, the highest genome recovery fraction, and the lowest number of contigs (Table S5).

We find that the *de fere novo* assembly using short reads performs better than *de novo* assembly using short reads alone. Comparison of *de fere novo* to hybrid assembly allows assessment of *de fere novo* accuracy, and indicates that *de fere novo* can recover the rDNA sequence correctly placed in their genomic context, with a low error rate.

### Case Study: Closing the assembly of *S. aureus* UAMS-1

*Staphylococcus aureus* UAMS-1 is a well-characterized, USA200 lineage, methicillin-sensitive strain isolated from an osteomyelitis patient. The published genome was sequenced using Illumina MiSeq (300bp reads), and the assembly refined with GapFiller as part of the BugBuilder pipeline http://www.imperial.ac.uk/bioinformatics-data-science-group/resources/software/bugbuilder/. Currently, the genome assembly is represented by two scaffolds (JTJK00000000), with several repeated regions acknowledged in the annotations [40]. As the rDNA regions were not fully characterized in the annotations, we proposed that *de fere novo* assembly might resolve some of the problematic regions.

Using the same reference *S. aureus* MRSA252 [18] (BX571856.1) with riboSeed as was used in the original assembly, *de fere novo* assembly correctly bridged gaps corresponding to three of the five rDNAs in the reference genome (Table 3B). Furthermore, *de fere novo* assembly bridged two contigs that were syntenic with the ends of the scaffolds in the published assembly, indicating that the regions resolved by riboSeed could improve closure of the genome.

We modified the BugBuilder pipeline (https://github.com/nickp60/BugBuilder) used in the published assembly to incorporate pseudocontigs from riboSeed. Further, we compared the performance of Pilon, GapFiller, and no finishing software with both the *de fere novo* and *de novo* assemblies (see Table S6). All assemblies resulted in a single scaffold (updates to BugBuilder and many of its dependencies prevented exact recapitulation of the published assembly), but scaffolds varied in length, number of ambiguous bases, and resolution of rDNA repeats. In all cases, riboSeed’s *de fere novo* assemblies resulted in more rDNA regions being resolved. In this case, riboSeed was able to assist in bringing an existing high-quality scaffold to near closure.

### Benchmarking against GAGE-B Datasets

We used the Genome Assembly Gold-standard Evaluation for Bacteria (GAGE-B) datasets [27] to assess the performance of riboSeed against a set of well-characterized assemblies. These datasets represent a broad range of challenges; low GC content and tandem rDNA repeats prove challenging to the riboSeed procedure.

*Mycobacterium abscessus* has only a single rDNA operon and does not suffer from the issue of rDNA repeats, so was excluded from this analysis. We also excluded the *B. cereus* VD118 HiSeq dataset, as metagenomic analysis revealed likely contamination (see Figure S5 and supplementary data).

When the reference used in the GAGE-B study was also the sequenced strain (e.g. *Rhodobacter sphaeroides* and *Bacillus cereus*), we chose an alternate reference, as using the original reference could provide an unfair advantage to riboSeed. The GAGE-B datasets include both raw and trimmed reads; in all cases, the trimmed reads were used. Results are shown in Table 3C.

Compared to *de novo* assembly, the *de fere novo* approach improved the majority of assemblies. In the case of the *S. aureus* and *R. sphaeroides* datasets, particular difficulty was encountered for all references tested. In the case of *Bacteroides fragilis*, the entropy plot (Figure S4.3) shows that sequence variability on the 5’ end of the operon is much lower within the genome compared to many of the other within-genome figures, possibly contributing to misassembly.

## Discussion

We demonstrate that regions flanking equivalent rDNAs from related strains show a high degree of conservation in related organisms. This allows us to correctly place rDNAs within a newly sequenced isolate, even in the absence of the resolution that would be provided by long read sequencing. Comparing the regions flanking rDNAs within a single genome, we observed that when considering sufficiently large flanking regions, flanking sequences show enough variability to differentiate each instance of the rDNAs. Taken together with the cross-taxon homology, this allows inference of the location (i.e. the flanking regions) of rDNAs, and the variability of these flanking regions within a genome enables unique identification of reads likely belonging to a specific cluster.

The extent of sequence similarity between the sequenced isolate and the reference influences *de fere novo* assembly. If fewer than 80% of reads map to the reference, resulting pseudocontigs are treated as “untrusted” contigs by SPAdes to prevent spurious joining of contigs. Figure 5 shows that although one should preferentially use the closest complete reference available for optimal results, the subassembly method is robust against moderate discrepancies between the reference and sequenced isolate’s flanking regions.

Strains possessing a single rDNA (such as *Mycobacterium abscessus*) do not suffer from repeated region assembly issues. Similarly, the rRNA coding regions in some taxa (such as *Thermus thermophilus* or *Leptospira interrogans*, see Supplementary section “Atypical rDNA operon structure”, Figure S6) are not organized into operons. Such genomes do not require correction with riboSeed.

The method of constructing pseudocontigs implemented by riboSeed relies on having a relevant reference sequence, where the rDNA regions act as “bait”, fishing for reads that likely map specifically to that region. This makes riboSeed a valuable tool for assembly or reassembly of bacterial or archaeal strains (Table S7) for which such a reference is available, but application to community ecology where one may be sequencing novel organisms from unsequenced genera will be limited by the requirement for such a reference genome. Although we show this “baiting” method to be an effective way to partition the reads, a more robust method may be to use a probabilistic representation of equivalent rDNA regions for the sequenced taxon. By using a database of sequence profiles (e.g. hidden Markov Models) from homologous rDNAs in a taxon, the step of choosing a single most appropriate reference might be circumvented. For datasets where the choice of reference determines riboSeed’s effectiveness, a probabilistic approach may improve performance.

Several checks are implemented after subassembly to ensure that resulting pseudocontigs are fit for inclusion in the next round in the next mapping/subassembly iteration or the final *de fere novo* assembly. If a subassembly’s longest contig is greater than 3x the particular pseudocontig length or shorter than 6kb (a conservative minimum length of a 16S, 23S, and 5S operon), this is taken to be a sign of poor parameter choice so the user is warned, and by default no further seedings will occur to avoid spurious assembly. Such an outcome can be indicative of any of several factors: improper clustering of operons; insufficient or extraneous flanking sequence; sub-optimal mapping; inappropriate choice of k-mer length for subassembly; inappropriate reference; or other issues. If this occurs, we recommend testing the assembly with different k-mers, changing the flanking length, or trying alternative reference genomes. Mapping depth of the rDNA regions is also reported for each iteration; a marked decrease in mapping depth may also be indicative of problems.

Many published genome finishing tools and approaches offer improvements when applied to suitable datasets, but none (including the approach presented in this paper) is able in isolation to resolve all bacterial genome assembly issues. One constraint on the performance of riboSeed is the quality of rRNA annotations in reference strains. Although it is impossible to concretely confirm *in silico*, we (and others [29]) have found several reference genomes during the course of this study that we suspect have collapsed rDNA repeats. We recommend using a tool such as 16Stimator [35] or rrnDB [42] to estimate number of 16S (and therefore rDNAs) prior to assembly, or `stack` to assess mapping depths after running `seed`.

As riboSeed relies on de Bruijn graph assembly, the results can be affected by assembler parameters. Care should be taken to explore appropriate settings, particularly in regard to read trimming approach, range of *k*-mers, and error correction schemes.

One difficulty in determining the accuracy of rDNA counts in reference genome occurs because genome sequences are often released without publishing the reads used to produce the genome assembly. This practice is a major hindrance when attempting to perform coverage-based quality assessment, such as to infer the likelihood of collapsed rDNAs. While data transparency is expected for gene expression studies, that stance has not been universally adopted when publishing whole-genome sequencing results. To ensure the highest quality assemblies from historical data, we strongly recommend that researchers share raw reads.

## Conclusions

Demonstration that rDNA flanking regions are conserved across taxa and that flanking regions of sufficient length are distinct within a genome allowed for the development of riboSeed, a *de fere novo* assembly method. riboSeed utilizes rDNA flanking regions to act as barcodes for repeated rDNAs, allowing the assembler to correctly place and orient the rDNA. *de fere novo* assembly can improve the assembly by bridging across ribosomal regions, and, in cases where rDNA repeats would otherwise result in incomplete scaffolding, can result in closure of a draft genome when used in conjunction with existing polishing tools. Although riboSeed is far from a silver bullet to provide perfect assemblies from short read technology, it shows the utility of using genomic reference data and mixed assembly approaches to overcome algorithmic obstacles. This approach to resolving rDNA repeats may allow further insight to be gained from large public repositories of short read sequencing data, such as SRA, and when used in conjunction with other genome finishing techniques, provides an avenue towards genome closure.

## List of abbreviations

rDNA: DNA region coding for ribosomal RNA operon
rRNA: ribosomal RNA
SRA: Sequence Read Archive
ENA: European Nucleotide Archive
IG: intergenic
GAGE-B: Genome Assembly Gold-standard Evaluation for Bacteria

## Competing interests

The authors declare that they have no competing interests

## Funding

The work was funded through a joint studentship between The James Hutton Institute, Dundee, Scotland, and the National University of Ireland, Galway, Ireland.

## Authors’ contributions

NRW wrote all the bugs.

## Acknowledgements

We thank Anton Korobeynikov for his recommendations on optimizing SPAdes. Yoann Augagneur, Shaun Brinsmade, and Mohamed Sassi graciously provided access to the *S. aureus* UAMS-1 genome sequencing data. Additional computational resources were provided by CLIMB [14]. We thank the Bioconda (https://bioconda.github.io/) community for their support.

## References

[1] Can Alkan, Saba Sajjadian, and Evan E Eichler. Limitations of next-generation genome sequence assembly. Nature Methods, 8(1), 2011.

[2] Anton Bankevich, Sergey Nurk, Dmitry Antipov, Alexey A Gurevich, Mikhail Dvorkin, Alexander S Kulikov, Valery M Lesin, Sergey I Nikolenko, Son Pham, Andrey D Prjibelski, Alexey V Pyshkin, Alexander V Sirotkin, Nikolay Vyahhi, Glenn Tesler, Max A Alekseyev, and Pavel A Pevzner. SPAdes: A New Genome Assembly Algorithm and Its Applications to Single-Cell Sequencing. Journal of Computational Biology, 19(5):455–477, 2012.

[3] Marten Boetzer and Walter Pirovano. Toward almost closed genomes with GapFiller. Genome Biology, 13(6), 2012.

[4] Balázs Brankovics, Hao Zhang, Anne D. van Diepeningen, Theo A. J. van der Lee, Cees Waalwijk, and G. Sybren de Hoog. GRAbB: Selective Assembly of Genomic Regions, a New Niche for Genomic Research. PLOS Computational Biology, 12(6):e1004753, 2016.

[5] Carlo P. J. M. Brouwer, Thuy Duong Vu, Miaomiao Zhou, Gianluigi Cardinali, Mick M. Welling, Nathalie van de Wiele, and Vincent Robert. Current Opportunities and Challenges of Next Generation Sequencing (NGS) of DNA; Determining Health and Disease. British Biotechnology Journal, 13(4), 2016.

[6] Rebecca J. Case, Yan Boucher, Ingela Dahllöf, Carola Holmström, W. Ford Doolittle, and Staffan Kjelleberg. Use of 16S rRNA and rpoB Genes as Molecular Markers for Microbial Ecology Studies. Applied and Environmental Microbiology, 73(1):278–288, 2007.

[7] Jill E Clarridge III. Impact of 16S rRNA gene sequence analysis for identification of bacteria on clinical microbiology and infectious diseases. Clinical Microbiology Reviews, 17(4):840–62, table of contents, 2004.

[8] Tom Coenye and Peter Vandamme. Intragenomic heterogeneity between multiple 16S ribosomal RNA operons in sequenced bacterial genomes. FEMS Microbiology Letters, 228:45–49, 2003.

[9] Aaron Darling, Andrew Tritt, Jonathan A. Eisen, and Marc T. Facciotti. Mauve Assembly Metrics. Bioinformatics Advance Access, 2011.

[10] Aaron C.E. Darling, Bob Mau, Frederick R. Blattner, and Nicole T. Perna. Mauve: Multiple Alignment of Conserved Genomic Sequence With Rearrangements. Genome Research, 14(7):1394–1403, 2004.

[11] Aarti Desai, Veer Singh Marwah, Akshay Yadav, Vineet Jha, Kishor Dhaygude, Ujwala Bangar, Vivek Kulkarni, and Abhay Jere. Identification of optimum sequencing depth especially for de novo genome assembly of small genomes using next generation sequencing data. PloS ONE, 8(4), 2013.

[12] Marco Galardini, Emanuele G Biondi, Marco Bazzicalupo, and Alessio Mengoni. CONTIGuator: a bacterial genomes finishing tool for structural insights on draft genomes. Source Code for Biology and Medicine, 6(11), 2011.

[13] Johan Goris, Konstantinos T Konstantinidis, Joel A Klappenbach, Tom Coenye, Peter Vandamme, and James M Tiedje. DNA–DNA hybridization values and their relationship to whole-genome sequence similarities. International Journal of Systematic and Evolutionary Microbiology, 57:81–91, 2007.

[14] Martyn Guest, Joel Southgate, Matthew Ismail, Marius Bakke, Radoslaw Poplawski, Thomas R. Connor, Nicholas J. Loman, Simon Elwood Thompson, Simon Thompson, Matthew J. Bull, Christine Kitchen, Andy Smith, Emily Richardson, Samuel K. Sheppard, and Mark J. Pallen. CLIMB (the Cloud Infrastructure for Microbial Bioinformatics): an online resource for the medical microbiology community. Microbial Genomics, 2(9), sep 2016.

[15] Dieval Guizelini, Roberto T. Raittz, Leonardo M. Cruz, Emanuel M. Souza, Maria B. R. Steffens, and Fabio O. Pedrosa. GFinisher: a new strategy to refine and finish bacterial genome assemblies. Nature Scientific Reports, 6, 2016.

[16] Alexey Gurevich, Vladislav Saveliev, Nikolay Vyahhi, and Glenn Tesler. QUAST: quality assessment tool for genome assemblies. Bioinformatics, 29(8):1072–1075, apr 2013.

[17] Tetsuya Hayashi, Kozo Makino, Makoto Ohnishi, Ken Kurokawa, Kazuo Ishii, Katsushi Yokoyama, Chang-Gyun Han, Eiichi Ohtsubo, Keisuke Nakayama, Takahiro Murata, Masashi Tanaka, Toru Tobe, Tetsuya Iida, Hideto Takami, Takeshi Honda, Chihiro Sasakawa, Naotake Ogasawara, Teruo Yasunaga, Satoru Kuhara, Tadayoshi Shiba, Masahira Hattori, and Hideo Shinagawa. Complete Genome Sequence of Enterohemorrhagic Eschelichia coli O157:H7 and Genomic Comparison with a Laboratory Strain K-12. DNA Research, 8:11–22, 2001.

[18] Matthew T. G. Holden, Edward J. Feil, Jodi A. Lindsay, Sharon J. Peacock, Nicholas P. J. Day, Mark C. Enright, Tim J. Foster, Catrin E. Moore, Laurence Hurst, Rebecca Atkin, Andrew Barron, Nathalie Bason, Stephen D. Bentley, Carol Chillingworth, Tracey Chillingworth, Carol Churcher, Louise Clark, Craig Corton, Ann Cronin, Jon Doggett, Linda Dowd, Theresa Feltwell, Zahra Hance, Barbara Harris, Heidi Hauser, Simon Holroyd, Kay Jagels, Keith D. James, Nicola Lennard, Alexandra Line, Rebecca Mayes, Sharon Moule, Karen Mungall, Douglas Ormond, Michael A. Quail, Ester Rabbinowitsch, Kim Rutherford, Mandy Sanders, Sarah Sharp, Mark Simmonds, Kim Stevens, Sally Whitehead, Bart G. Barrell, Brian G. Spratt, and Julian Parkhill. Complete genomes of two clinical Staphylococcus aureus strains: evidence for the rapid evolution of virulence and drug resistance. Proceedings of the National Academy of Sciences of the United States of America, 101(26):9786–91, jun 2004.

[19] Xuesong Hu, Jianying Yuan, Yujian Shi, Jianliang Lu, Binghang Liu, Zhenyu Li, Yanxiang Chen, Desheng Mu, Hao Zhang, Nan Li, Zhen Yue, Fan Bai, Heng Li, and Wei Fan. pIRS: Profile-based Illumina pair-end reads simulator. Bioinformatics, 28(11):1533–1535, jun 2012.

[20] Kazutaka Katoh, Kazuharu Misawa, Keiichi Kuma, and Takashi Miyata. MAFFT: a novel method for rapid multiple sequence alignment based on fast Fourier transform. Nucleic Acids Research, 30(14):3059–3066, jul 2002.

[21] Paul A Kitts, Deanna M Church, Françoise Thibaud-Nissen, Jinna Choi, Vichet Hem, Victor Sapojnikov, Robert G Smith, Tatiana Tatusova, Charlie Xiang, Andrey Zherikov, Michael DiCuccio, Terence D Murphy, Kim D Pruitt, and Avi Kimchi. Assembly: a resource for assembled genomes at NCBI. Nucleic acids research, 44(D1):D73–80, jan 2016.

[22] Yuichi Kodama, Martin Shumway, and Rasko Leinonen. The sequence read archive: explosive growth of sequencing data. Nucleic Acids Research, 40(Database issue):D54–56, jan 2012.

[23] Heng Li, Bob Handsaker, Alec Wysoker, Tim Fennell, Jue Ruan, Nils Homer, Gabor Marth, Goncalo Abecasis, Richard Durbin, and 1000 Genome Project Data Processing Subgroup. The Sequence Alignment/Map format and SAMtools. Bioinformatics, 25(16):2078–9, aug 2009.

[24] Pinglei Liu, Peng Li, Xiaofei Jiang, Dexi Bi, Yingzhou Xie, Cui Tai, Zixin Deng, Kumar Rajakumar, and Hong-Yu Ou. Complete genome sequence of Klebsiella pneumoniae subsp. pneumoniae HS11286, a multidrug-resistant strain isolated from human sputum. Journal of Bacteriology, 194(7):1841–1842, apr 2012.

[25] Oksana Lukjancenko, Trudy M. Wassenaar, and David W. Ussery. Comparison of 61 Sequenced Escherichia coli Genomes. Microbial Ecology, 60, 2010.

[26] Ruibang Luo, Binghang Liu, Yinlong Xie, Zhenyu Li, Weihua Huang, Jianying Yuan, Guangzhu He, Yanxiang Chen, Qi Pan, Yunjie Liu, Jingbo Tang, Gengxiong Wu, Hao Zhang, Yujian Shi, Yong Liu, Chang Yu, Bo Wang, Yao Lu, Changlei Han, David W Cheung, Siu-Ming Yiu, Shaoliang Peng, Zhu Xiaoqian, Guangming Liu, Xiangke Liao, Yingrui Li, Huanming Yang, Jian Wang, Tak-Wah Lam, and Jun Wang. SOAPdenovo2: an empirically improved memory-efficient short-read de novo assembler. GigaScience, 1(1):18, 2012.

[27] Tanja Magoc, Stephan Pabinger, Stefan Canzar, Xinyue Liu, Qi Su, Daniela Puiu, Luke J. Tallon, and Steven L. Salzberg. GAGE-B: an evaluation of genome assemblers for bacterial organisms. Bioinformatics, 29(14):1718–1725, 2013.

[28] Diego CB Mariano, Felipe L Pereira, Preetam Ghosh, Debmalya Barh, Henrique CP Figueiredo, Artur Silva, Rommel TJ Ramos, and Vasco AC Azevedo. MapRepeat: an approach for effective assembly of repetitive regions in prokaryotic genomes. Bioinformatician, 11(6):276–279, 2015.

[29] Diego César Batista Mariano, Thiago De Jesus Sousa, Felipe Luiz Pereira, Flávia Aburjaile, Debmalya Barh, Flávia Rocha, Anne Cybelle Pinto, Syed Shah Hassan, Tessália Diniz, Luerce Saraiva, Fernanda Alves Dorella, Alex Fiorini De Carvalho, Carlos Augusto Gomes Leal, Henrique César, Pereira Figueiredo, Artur Silva, Rommel Thiago, Jucá Ramos, Vasco Ariston, and Carvalho Azevedo. Whole-genome optical mapping reveals a misassembly between two rRNA operons of Corynebacterium pseudotuberculosis strain 1002. BMC Genomics, 17, 2016.

[30] Mari Miyamoto, Daisuke Motooka, Kazuyoshi Gotoh, Takamasa Imai, Kazutoshi Yoshitake, Naohisa Goto, Tetsuya Iida, Teruo Yasunaga, Toshihiro Horii, Kazuharu Arakawa, Masahiro Kasahara, and Shota Nakamura. Performance comparison of second-and third-generation sequencers using a bacterial genome with two chromosomes. BMC Genomics, 15, 2014.

[31] Claudia Moreno, Jaime Romero, and Romilio T. Espejo. Polymorphism in repeated 16S rRNA genes is a common property of type strains and environmental isolates of the genus Vibrio. Microbiology, 148:1233–1239, 2002.

[32] Francesca Nadalin, Francesco Vezzi, and Alberto Policriti. GapFiller: a de novo assembly approach to fill the gap within paired reads. BMC Bioinformatics, 13:12–14, 2012.

[33] Niranjan Nagarajan, Christopher Cook, MariaPia Di Bonaventura, Hong Ge, Allen Richards, Kimberly A Bishop-Lilly, Robert Desalle, Timothy D Read, and Mihai Pop. Finishing genomes with limited resources: lessons from an ensemble of microbial genomes. BMC Genomics, 11(1):242, 2010.

[34] Makoto Ohnishi, Takahiro Muratal, Keisuke Nakayama, Satoru Kuhara, Masahiro Hattori, Ken Kurokawa, Teruo Yasunaga, K Atsushi Yokoyamas, Kozo Makinos, Hideo Shinagawa, and Tetsuya Hayashi. Comparative Analysis of the Whole Set of rRNA Operons Between an Enterohemorrhagic Escherichia coli 0157:H7 Sakai Strain and an Escherichia coli K-12 Strain MG1655. Systematic and Applied Microbiology, 23:315–324, 2000.

[35] Matthew Perisin, Madlen Vetter, Jack A Gilbert, and Joy Bergelson. 16Stimator: statistical estimation of ribosomal gene copy numbers from draft genome assemblies. The ISME Journal, 10(4):1020–1024, apr 2016.

[36] Vitor C Piro, Helisson Faoro, Vinicius A Weiss, Maria Br Steffens, Fabio O Pedrosa, Emanuel M Souza, and Roberto T Raittz. FGAP: an automated gap closing tool. BMC Research Notes, 7, 2014.

[37] Aaron R. Quinlan and Ira M. Hall. BEDTools: a flexible suite of utilities for comparing genomic features. Bioinformatics, 26(6):841–84210, 2010.

[38] Peter Rice, Ian Longden, and Alan Bleasby. EMBOSS: the European Molecular Biology Open Software Suite. Trends in Genetics, 16(6):276–7, jun 2000.

[39] Fatemeh Sanjar, S L Rajasekhar Karna, Tsute Chen, Ping Chen, Johnathan J Abercrombie, and Kai P Leung. Whole-Genome Sequence of Multidrug-Resistant Pseudomonas aeruginosa Strain BAMCPA07-48, Isolated from a Combat Injury Wound. Genome Announcements, 4(4), 2016.

[40] Mohamed Sassi, Deepak Sharma, Shaun R Brinsmade, Brice Felden, and Yoann Augagneur. Genome Sequence of the Clinical Isolate Staphylococcus aureus subsp. aureus Strain UAMS-1. Genome Announcements, 3(1), 2015.

[41] Armin O. Schmitt and Hanspeter Herzel. Estimating the Entropy of DNA Sequences Introduction: Order and Disorder of Sequences. Journal of Theoretical Biology, 1888:369–377, 1997.

[42] Steven F. Stoddard, Byron J. Smith, Robert Hein, Benjamin R. K. Roller, and Thomas M. Schmidt. rrnDB: improved tools for interpreting rRNA gene abundance in bacteria and archaea and a new foundation for future development. Nucleic Acids Research, 43(D1), 2014.

[43] Todd J Treangen, Brian D Ondov, Sergey Koren, and Adam M Phillippy. The Harvest suite for rapid core-genome alignment and visualization of thousands of intraspecific microbial genomes. Genome Biology, 15(524), 2014.

[44] Todd J Treangen and Steven L Salzberg. Repetitive DNA and next-generation sequencing: computational challenges and solutions. Nature Reviews Genetics, 13(1):36–46, 2011.

[45] Isheng J Tsai, Thomas D Otto, and Matthew Berriman. Improving draft assemblies by iterative mapping and assembly of short reads to eliminate gaps. Genome Biology, 11, 2010.

[46] Sagar M Utturkar, Dawn M Klingeman, Miriam L Land, Christopher W Schadt, Mitchel J Doktycz, Dale A Pelletier, and Steven D Brown. Evaluation and validation of de novo and hybrid assembly techniques to derive high-quality genome sequences. Bioinformatics, 30(19):2709–2716, 2014.

[47] Benjamin P Vandervalk, Chen Yang, Zhuyi Xue, Karthika Raghavan, Justin Chu, Hamid Mohamadi, Shaun D Jackman, Readman Chiu, René L Warren, and Inanç Birol. Konnector v2.0: pseudo-long reads from paired-end sequencing data. BMC Medical Genomics, 8:2–5, 2015.

[48] Tomáš Větrovský and Petr Baldrian. The Variability of the 16S rRNA Gene in Bacterial Genomes and Its Consequences for Bacterial Community Analyses. PLoS ONE, 8(2), 2013.

[49] Bruce J. Walker, Thomas Abeel, Terrance Shea, Margaret Priest, Amr Abouelliel, Sharadha Sakthikumar, Christina A. Cuomo, Qiandong Zeng, Jennifer Wortman, Sarah K. Young, and Ashlee M. Earl. Pilon: An Integrated Tool for Comprehensive Microbial Variant Detection and Genome Assembly Improvement. PLoS ONE, 9(11):e112963, nov 2014.

[50] Dapeng Wang, Jiayue Xu, and Jun Yu. KGCAK: a K-mer based database for genome-wide phylogeny and complexity evaluation. Biology Direct, 10(53), 2015.

[51] Nicholas R Waters, Florence Abram, Ashleigh Holmes, Fiona Brennan, and Leighton Pritchard. riboSeed 0.4.35. Zenodo, 1(10.5281/zenodo.1037965), 2017.

[52] William G. Weisburg, Susan M. Barns, Dale A. Pelletier, and David J. Lane.16 Sribosomal DNA amplification for phylogenetic study. Journal of Bacteriology, 173(2):697–703, 1991.

[53] Nava Whiteford, Niall Haslam, Gerald Weber, Adam Prugel-Bennett, Jonathan W. Essex, Peter L. Roach, Mark Bradley, and Cameron Neylon. An analysis of the feasibility of short read sequencing. Nucleic Acids Research, 33(19), 2005.

[54] Carl R Woese, Otto Kandlert, and Mark L Wheelis. Towards a natural system of organisms: Proposal for the domains Archaea, Bacteria, and Eucarya. Proceedings of the National Academy of Sciences of the United States of America, 87:4576–4579, 1990.

[55] Derrick E Wood and Steven L Salzberg. Kraken: ultrafast metagenomic sequence classification using exact alignments. Genome Biology, 15(R46), 2014.

[56] Xing Yang, Daniel Medvin, Giri Narasimhan, Deborah Yoder-Himes, and Stephen Lory. CloG: a pipeline for closing gaps in a draft assembly using short reads. 2011 IEEE 1st International Conference on Computational Advances in Bio and Medical Sciences, pages 202–207, 2011.

